# Fault-tolerant pedigree reconstruction from pairwise kinship relations

**DOI:** 10.1101/2025.08.21.671608

**Authors:** Edward C. Huang, Kevin A. Li, Vagheesh M. Narasimhan

## Abstract

Pedigrees reconstructed from biologically related ancient genomes have revealed many insights into (pre)history. To our knowledge, all reported ancient pedigrees have been manually reconstructed, as existing pedigree reconstruction methods are ill-suited for the quality and nature of ancient DNA data. Here, we introduce `repare`, an open-source software method to automatically reconstruct pedigrees from inferred pairwise kinship relations, which are readily obtainable from ancient genomes. This method reconstructs pedigrees by iteratively incorporating pairwise kinship relations into a set of candidate pedigrees, with pruning and sampling to reduce its search space. It optionally considers supporting information such as haplogroups and skeletal age-at-death estimates. We evaluate this method on a variety of simulated pedigrees with varying error rates and missingness. We also use this method to reconstruct several published pedigrees that were originally manually reconstructed; for one, we present a potential alternative topology. `repare` optionally incorporates user-inferred pedigree constraints, enabling “human-in-the-loop” reconstruction workflows. Especially when used with these user-inferred constraints, we find that `repare` represents a powerful and flexible tool for ancient pedigree reconstruction.

## 1 Introduction

Advances in ancient DNA (aDNA) sequencing methods have enabled the sampling of ancient genomes from multiple individuals interred at the same archaeological site. Notably, a set of biologically related ancient genomes can be arranged into a pedigree, which in turn allows for analysis of inheritance patterns, population structure, and other social and biological trends through time. There exist many pedigree reconstruction methods built primarily for modern genetic samples [1–8]; these methods have seen little, if any, usage with ancient datasets. This is at least partially due to the unique nature of aDNA data: calls are often “pseudo-haploid” due to low sequence coverage [9], contamination and damage rates are elevated [10], exact birth years are unknowable with current dating techniques [11], and pedigrees are rarely completely sampled and can span many generations. These limitations break many of the assumptions of existing pedigree reconstruction methods and make it exceedingly difficult to reconstruct ancient pedigrees using methods not purpose-built for aDNA data.

As an example, `PRIMUS` is a state-of-the-art (modern) pedigree reconstruction method that utilizes identity-by-descent (IBD) proportion estimates [8]. However, `PRIMUS` is not designed to reconstruct consanguineous pedigrees, which can be common in ancient contexts. In addition, accurate IBD inference in ancient samples requires at least 1x mean coverage on the 1240k single nucleotide polymorphism capture panel [12], which precludes inclusion of many ancient samples. For example, the Hazleton North site has 7 individuals with less than 1x mean 1240k coverage out of 35 reported individuals [13], and the Gurgy ‘les Noisats’ site has 35 individuals with less than 1x mean 1240k coverage out of 94 reported individuals [14]. For these reasons, to our knowledge, all reported ancient pedigrees have been primarily manually reconstructed [13–26].

Despite aDNA’s relatively lower quality, it is feasible to infer pairwise kinship relations from ancient samples with as low as 0.05x average sequence coverage, albeit at non-negligible error rates [27–36]. As such, many ancient pedigrees have been manually reconstructed from inferred kinship relations along with supporting information including haplogroups and archaeological features such as skeletal age-at-death [13–26]. These ancient pedigrees have yielded numerous insights into burial practices, kinship-based social structure, and the spread of disease throughout (pre)history.

Although the reconstruction of ancient pedigrees has furthered our understanding of the past, the manual pedigree reconstruction process remains an extremely time-consuming endeavor. Here, we introduce `repare`, an open-source software method that automates pedigree reconstruction using information readily available from aDNA data. `repare` iteratively builds up a set of candidate pedigrees from inferred pairwise degree-level kinship relations, and uses optional supporting information including haplogroups, runs of homozygosity (ROH), skeletal age-at-death estimates, and manual relation constraints to prune invalid pedigrees. Since kinship inference from aDNA is imperfect, especially for lower-coverage samples, `repare` explicitly considers alternative possibilities for kinship relations when building pedigrees (Methods: Robustness to kinship inference errors). Using `repare`, we reconstruct pedigrees from a variety of simulated and published datasets, with a focus on aDNA data. We find that `repare` can accurately reconstruct both simulated and published pedigrees from imperfect kinship relations, offering a valuable tool for automatic pedigree reconstruction.

## 2 Methods

### 2.1 Iterative pedigree reconstruction

`repare`’s pedigree reconstruction algorithm iteratively incorporates pairwise kinship relations into a set of progressively expanding candidate pedigrees. At each iteration, `repare` considers a new degree-level kinship relation between two individuals. Inferred first- and second-degree relations are explicitly incorporated while considering potential alternative inferences (Methods: Robustness to kinship inference errors); third-degree relation information is used to help score pedigrees (Methods: Third-degree kinship relations). Input kinship relations are sorted by ascending degree. Within each degree, kinship relations are sorted by descending individual connectivity so that relations involving individuals with more total relations are reconstructed first. We hypothesize that this ordering can help constrain later kinship relations by first constructing dense subpedigrees.

Then, in each existing candidate pedigree, `repare` incorporates all possible exact kinship relations between the two individuals, barring relation type constraints (Methods: Robustness to kinship inference errors). Because each degree-level relationship can correspond to multiple exact relations, `repare` duplicates candidate pedigrees before incorporating new relations in order to represent each possible scenario. In addition, `repare` uses placeholder individuals to represent unsampled individuals in candidate pedigrees. For each input relation, users can also set constraints on the exact relation type, for example for first-degree relations where kinship relation inference methods can differentiate parental and full-sibling relations. `repare` does not restrict the number of kinship relations shared between individuals, enabling reconstruction of inbred relationships.

### 2.2 Pedigree pruning

Because degree-level kinship relations can be represented as multiple different relation configurations, the number of possible pedigrees increases exponentially with the number of algorithm iterations (i.e., the number of input kinship relations). At each algorithm iteration, supplementary individual-level genetic and archaeological data are used to prune the set of candidate pedigrees. All such data are optional inputs, except for genetic sex, which we use to assign parentage. These supplementary data can include haplogroups and skeletal age-at-death estimates, which we use to remove candidate pedigrees with invalid relations. If available, ROH data are also used to eliminate candidate pedigrees with inbred individuals that we know are not inbred. Finally, years-before-present data are used to eliminate candidate pedigrees with relations that are temporally improbable, for example based on archaeological layer information or significant carbon dating differences.

### 2.3 Robustness to kinship inference errors

Inference of kinship relations from aDNA data is not perfectly accurate, and performance tends to deteriorate with lower sequence coverage and more distant kinship. As such, most datasets of inferred kinship relations from aDNA contain multiple errors, in which case an iterative pedigree construction algorithm that treats input kinship relations as fixed cannot reconstruct the true pedigree. In addition, in many cases, as few as one incorrect kinship relation can make finding even one valid pedigree impossible. To address this limitation, we implement an “inconsistency” scoring system for pedigrees: `repare` explicitly considers alternative possibilities for each input kinship relation; then, in each resulting candidate pedigree, each conflict between a relation in a candidate pedigree and the input kinship data is considered one inconsistency. For example, `repare` will try changing an inferred first-degree relation to a second-degree relation if that better fits the reconstructed pedigree. Then, after all input relations are incorporated, the pedigree with the fewest inconsistencies against the input data is returned.

### 2.4 Third-degree kinship relations

`repare` explicitly incorporates first- and second-degree kinship relations into candidate pedigrees and uses third-degree kinship relations as a tiebreaker mechanism to differentiate between otherwise equally plausible pedigrees. We choose not to explicitly incorporate third-degree kinship relations because of their relatively higher inference error rates and the additional complexity involved with enumerating all possible third-degree exact kinship relation types; not only are third-degree relations more common in most pedigrees, there are more possible exact types of each third-degree relation. As such, we utilize the information from inferred third-degree kinship relations by introducing a separate “third-degree inconsistency” system that counts the number of extraneous third-degree kinship relations present in a candidate pedigree but not in the input kinship data. Since we do not explicitly incorporate third-degree kinship relations, we do not penalize third-degree relations present in the input data but not in a candidate pedigree. We use the third-degree inconsistency metric to differentiate candidate pedigrees with the same number of (first- and second-degree) inconsistencies.

### 2.5 Pedigree sampling

Even with pedigree pruning, we find that the number of candidate pedigrees grows far too quickly to enumerate every possible final pedigree. Therefore, we implement a sampling scheme to constrain the number of candidate pedigrees kept after each iteration. After incorporating each kinship relation into the set of candidate pedigrees, `repare` downsamples the candidate pedigree set to a fixed, user-defined number of pedigrees; we set the default value of this parameter to 1000. We downsample candidate pedigrees using a form of epsilon-greedy sampling [37], which takes the highest-scoring (greedy) action with probability 1 *− ε* and a random action with probability *ε*, adapted for sampling multiple actions. Let *n* be the maximum number of candidate pedigrees to keep after each algorithm iteration and let *ε* be the “batch epsilon-greedy” parameter. To avoid running conventional epsilon-greedy sampling *n* times (without replacement), after each algorithm iteration we simply define the new, downsampled candidate pedigree set in two parts: ignoring integer rounding, we deterministically select the (1 *− ε*)*n* candidate pedigrees with the fewest inconsistencies, and then uniformly sample *n −* (1 *− ε*)*n* pedigrees from the remaining set of candidate pedigrees. We set the default value of *ε* to 0.2, although we note that varying *ε* appears to have little effect on reconstruction performance (Results: Simulated pedigree reconstruction). We evaluate the runtime and memory usage of `repare` in Supplementary Note 3: Algorithm runtime and memory usage.

### 2.6 Pedigree reconstruction evaluation

To measure pedigree reconstruction performance, we record two metrics: relation F1 score and degree F1 score. We define the relation F1 score as the harmonic mean of precision and recall over the set of exact kinship relations (e.g., maternal grandparent-grandchild) in the ground-truth pedigree. Similarly, we define the degree F1 score as the harmonic mean of precision and recall over the set of degree-level kinship relations (e.g., first-degree) in the ground-truth pedigree. Both metrics consider only first- and second-degree relations between sampled (non-missing) individuals. While the relation F1 score evaluates reconstruction of the exact simulated pedigree, we hypothesize that many ancient pedigree reconstruction problems are underdetermined such that multiple final pedigrees could plausibly agree with the input kinship relations data. Therefore, we also record degree F1 scores, which helps evaluate a more general structure of a predicted pedigree.

## 3 Results

### 3.1 Simulated pedigree reconstruction

We assess `repare`’s performance in reconstructing a variety of simulated pedigrees. To do so, we implement a custom pedigree simulator that generates ground-truth pedigrees and their accompanying genetic and age-at-death data (Supplementary Note 1: Ancient pedigree simulation). We utilize this pedigree simulator to generate ground-truth pedigrees and their accompanying genetic and age-at-death data, approximating the data collected from aDNA. To evaluate `repare`’s performance under a range of data quality regimes, we vary the simulator’s *p*(mask node) parameter, which corresponds to pedigree missingness. We also vary its simulated sequence coverage parameter, which approximates kinship relation inference error rates based on those of `KIN` [34], a popular ancient kinship inference method, in a performance benchmark [38] (Figure 2). For each configuration of the *p*(mask node) and simulated sequence coverage parameters, we use the same 100 simulated ground-truth pedigrees for consistency. Then, we mask and corrupt the data of each pedigree corresponding to the experiment’s parameters before providing it as input to `repare`. In this experiment, we reconstruct pedigrees using the default parameters of 1000 for the maximum number of candidate pedigrees kept after each algorithm iteration and 0.2 for the “batch epsilon-greedy” parameter (Methods: Pedigree sampling).

**Figure 1.**
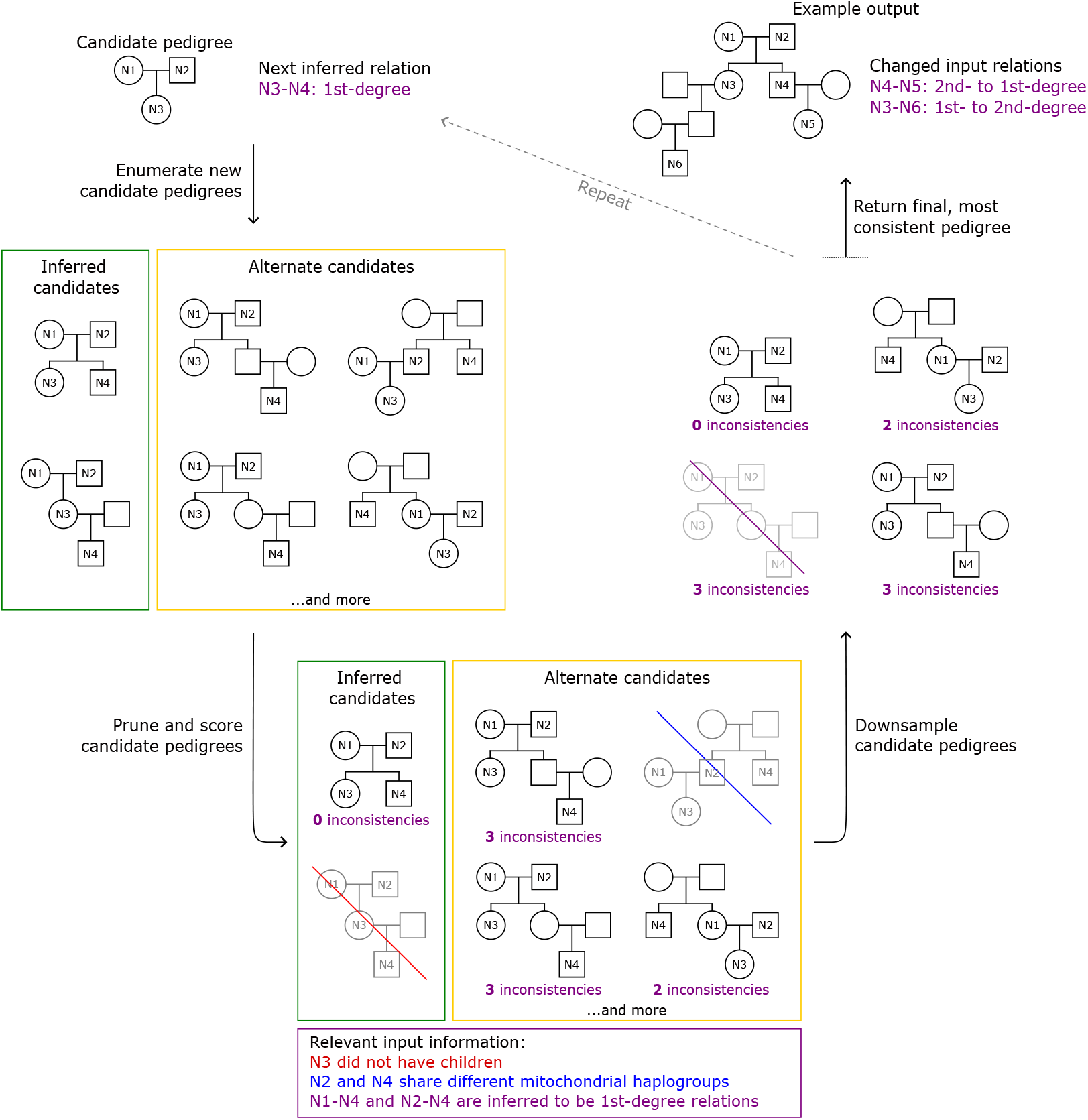
Diagram of `repare’s` pedigree reconstruction algorithm (Methods: Iterative pedigree reconstruction). At each algorithm iteration, an inferred kinship relation is incorporated into the current set of candidate pedigrees; alternative possibilities for the inferred relation are explicitly considered. The new set of candidate pedigrees is then pruned and downsampled before the next algorithm iteration.

**Figure 2.**
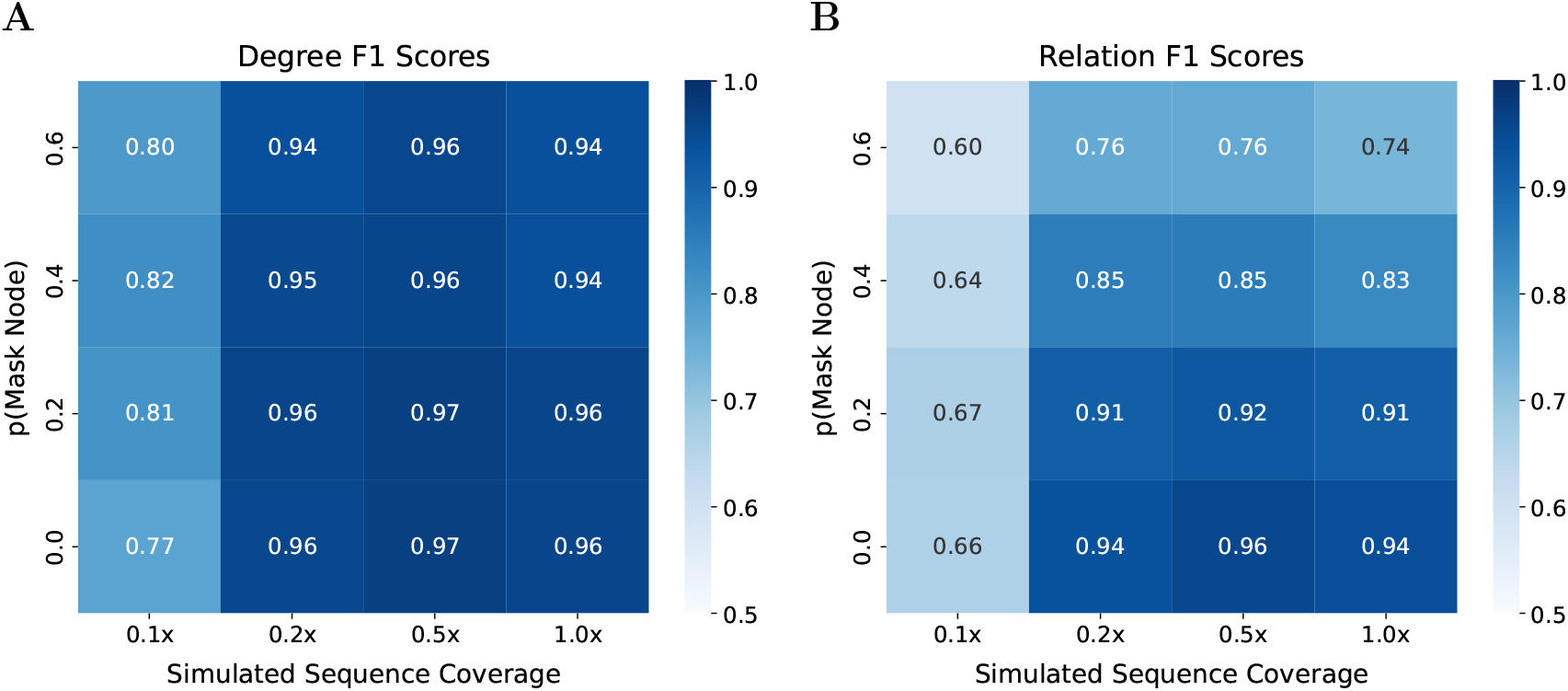
Pedigree reconstruction performance of `repare` on simulated data. We report reconstruction performance against ground-truth pedigrees using F1 scores, calculated over **(A)** degree-level kinship relations and **(B)** exact kinship relations (Methods: Pedigree reconstruction evaluation).

We measure reconstruction performance with degree and relation F1 scores (Methods: Pedigree reconstruction evaluation). In terms of both metrics, `repare` accurately reconstructs pedigrees, with relation F1 scores exceeding 0.9 and degree F1 scores exceeding 0.95, when data quality is high; in addition, `repare`’s degree F1 performance is more robust to decreases in data quality. We observe a steep decline in performance as simulated sequence coverage drops from 0.2x to 0.1x. We note that `repare` appears to perform worse at 1.0x simulated sequence coverage than at 0.5x simulated sequence coverage. A likely explanation for this phenomenon is that, in the benchmark experiment used to determine kinship error rates [38], `KIN` performs worse in some respects at 1.0x simulated sequence coverage than at 0.5x simulated sequence coverage. We further analyze pedigree characteristics and pedigree-level performance in Supplementary Note 2: Analysis of simulated pedigree reconstruction.

We also perform an experiment to quantify the effect of sampling parameters on reconstruction performance. Using the same 100 simulated ground-truth pedigrees as before, we evaluate `repare`’s performance while varying *ε*, the “batch epsilon-greedy” parameter, and the maximum number of candidate pedigrees kept after each algorithm iteration (Methods: Pedigree sampling) (Figure 3). To generate data from the 100 simulated pedigrees, we fix *p*(mask node) to 0.4 and the simulated coverage level to 0.5x. We find that varying *ε* appears to have a minimal impact on reconstruction performance. On the other hand, increasing the maximum number of candidate pedigrees kept after each algorithm iteration appears to slightly improve performance. To balance performance and runtime considerations, we set the default value of this “max candidate pedigrees” parameter to 1000.

**Figure 3.**
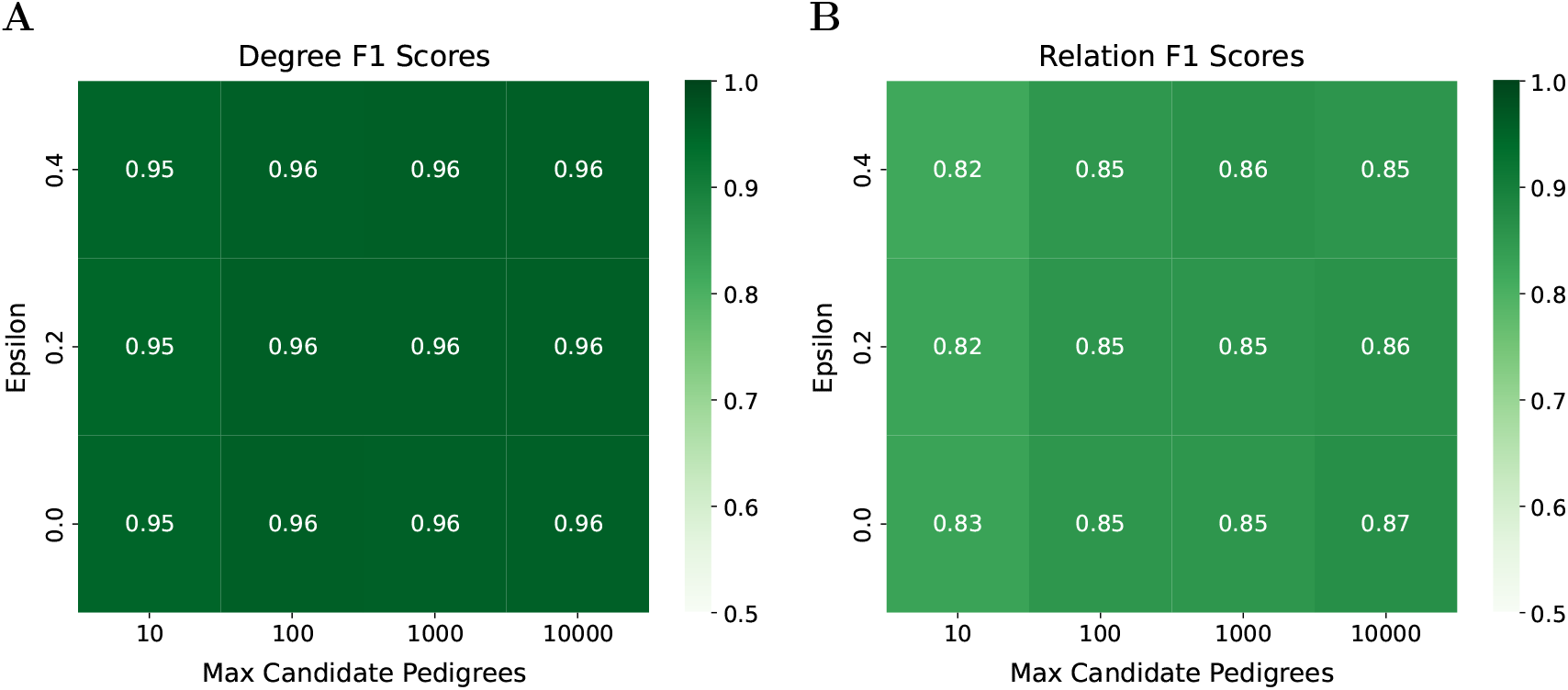
The effect of sampling parameters on `repare`’s pedigree reconstruction performance. We report reconstruction performance against simulated pedigrees using F1 scores, calculated over **(A)** degree-level kinship relations and **(B)** exact kinship relations (Methods: Pedigree reconstruction evaluation).

### 3.2 Published pedigree reconstruction

We also evaluate `repare`’s ability to reconstruct previously published ancient pedigrees. For each pedigree, we collect published kinship relation inference data and genetic and archaeological data including haplogroups, genetic sexes, skeletal age-at-death estimates, and ROH. We provide these data as input to `repare` and assess its pedigree reconstruction performance using the published pedigree as ground truth. In certain cases, we also incorporate additional contextual information from case-specific inferences made by the authors of the published papers; this is to illustrate `repare`’s potential as an assistive tool in “human-in-the-loop” pedigree reconstructions. We include only author inferences that independently distinguish between possible kinship relations and exclude those that use information *post hoc* from adjacent kinship relations. For example, if authors modify an inferred kinship relation to “fit” better with the adjacent individuals in a pedigree, we do not include this inference. We evaluate `repare`’s reconstructed pedigrees against the corresponding published pedigrees using relation F1 and degree F1 scores (Methods: Pedigree reconstruction evaluation). Some published pedigrees contain uncertain degree-level kinship relations where the authors do not infer an exact relation; these relations are often denoted with dotted lines in pedigree figures. Since `repare` only outputs pedigrees with exact relations, to calculate F1 scores over uncertain published relations, we consider a `repare`-reconstructed relation correct if it agrees with the degree of an uncertain relation and incorrect otherwise. Results for reconstructed pedigrees are summarized in Table 1.

**Table 1.**
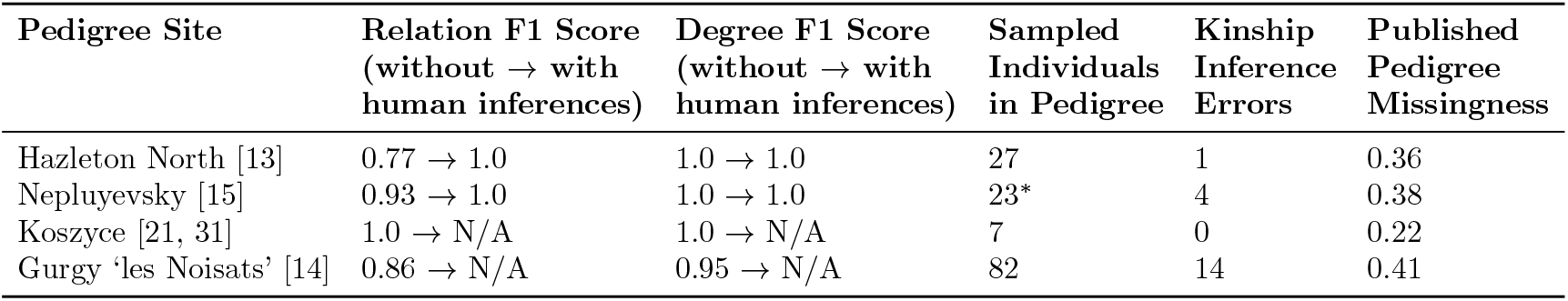
Results of `repare`’s reconstruction of published ancient pedigrees from their accompanying genetic and archaeological datasets. All pedigrees were reconstructed with default sampling parameters (Methods: Pedigree sampling). We report relation F1 and degree F1 scores (Methods: Pedigree reconstruction evaluation) to evaluate performance. When applicable, we report performance before and after providing `repare` with additional information from author inferences used to reconstruct the published pedigree. To provide context on reconstruction difficulty, we report the number of sampled individuals related to at least one other individual in the published pedigrees, the number of first- and second-degree kinship inference errors in the reported data with respect to the published pedigrees, and the missingness of each published pedigree. Published pedigree missingness is calculated by counting the number of plotted placeholder individuals, without considering uncertain relations, and dividing by the total number of individuals related to at least one other individual. ^*^The Nepluyevsky pedigree contains 23 individuals, but it is inferred by KIN to include a pair of identical twins, which repare treats as one individual.

#### 3.2.1 Hazleton North site

We assess `repare`’s performance in reconstructing a Neolithic pedigree from the Hazleton North site [13], using published genetic and archaeological data. The published pedigree was manually reconstructed using pairwise kinship relations and additional genetic and archaeological information including haplogroups, ROH, and skeletal age-at-death estimates. Degree-level kinship relations were inferred using relatedness coefficients (*r*), and first-degree relation types (parental or sibling) were inferred using IBD information. Using only this published data as input, `repare`’s reconstructed pedigree achieves a relation F1 score of 0.77 and a degree F1 score of 1.0 against the published pedigree. However, Fowler et al. [13] incorporated additional logical inferences to reconstruct their final published pedigree. In some cases, the directionality of parent-offspring kinship relations was determined through haplogroup rarity; for example, if a genetic male and a genetic female share a rare mitochondrial haplogroup as well as a parent-offspring kinship relation of unknown directionality, it is more likely that the female is the mother of the male and directly passed that haplogroup down. In another case, the final pedigree was selected over an alternative pedigree after manual inspection of the location of recombination breakpoints. When we include this additional information in `repare`’s inputs as kinship relation constraints, the reconstructed pedigree achieves perfect relation F1 and degree F1 scores of 1.0. However, we note that the reconstructed pedigree does not include SP4m as a connected individual, since SP4m’s closest inferred kinship relation is of the third degree. Overall, this result demonstrates `repare`’s effectiveness not only as a fully automatic pedigree reconstruction tool but also as a useful tool for iterative semi-automatic analyses.

#### 3.2.1 Nepluyevsky site

We also evaluate `repare`’s reconstruction of a Bronze Age pedigree from the Nepluyevsky site [15]. Similar to the Hazleton North pedigree, the published Nepluyevsky pedigree was manually reconstructed using pairwise kinship relations and supporting information including haplogroups and skeletal age-at-death estimates. Degree-level kinship relations and first-degree relation types were inferred using `KIN` [34]. Using this published data as input, `repare`’s reconstructed pedigree achieves a relation F1 score of 0.93 and a degree F1 score of 1.0 against the published pedigree. However, we note that `repare`’s reconstructed pedigree appears to represent a valid alternative pedigree for the data. There are a number of second-degree relations for which `repare` inferred a different relation type than in the published pedigree. For these relations, it is unclear how the final relation type was determined for the published pedigree. Therefore, we believe that `repare`’s reconstructed pedigree represents a plausible alternative pedigree given the available data. The published pedigree and original reconstructed pedigree are shown in Figure 4. Assuming each differing relation type was determined through author inferences, we encode this information into `repare`’s inputs as relation constraints, similar to our procedure for the Hazleton North pedigree (Results: Hazleton North site). When we do so, the reconstructed pedigree exactly matches the published pedigree.

**Figure 4.**
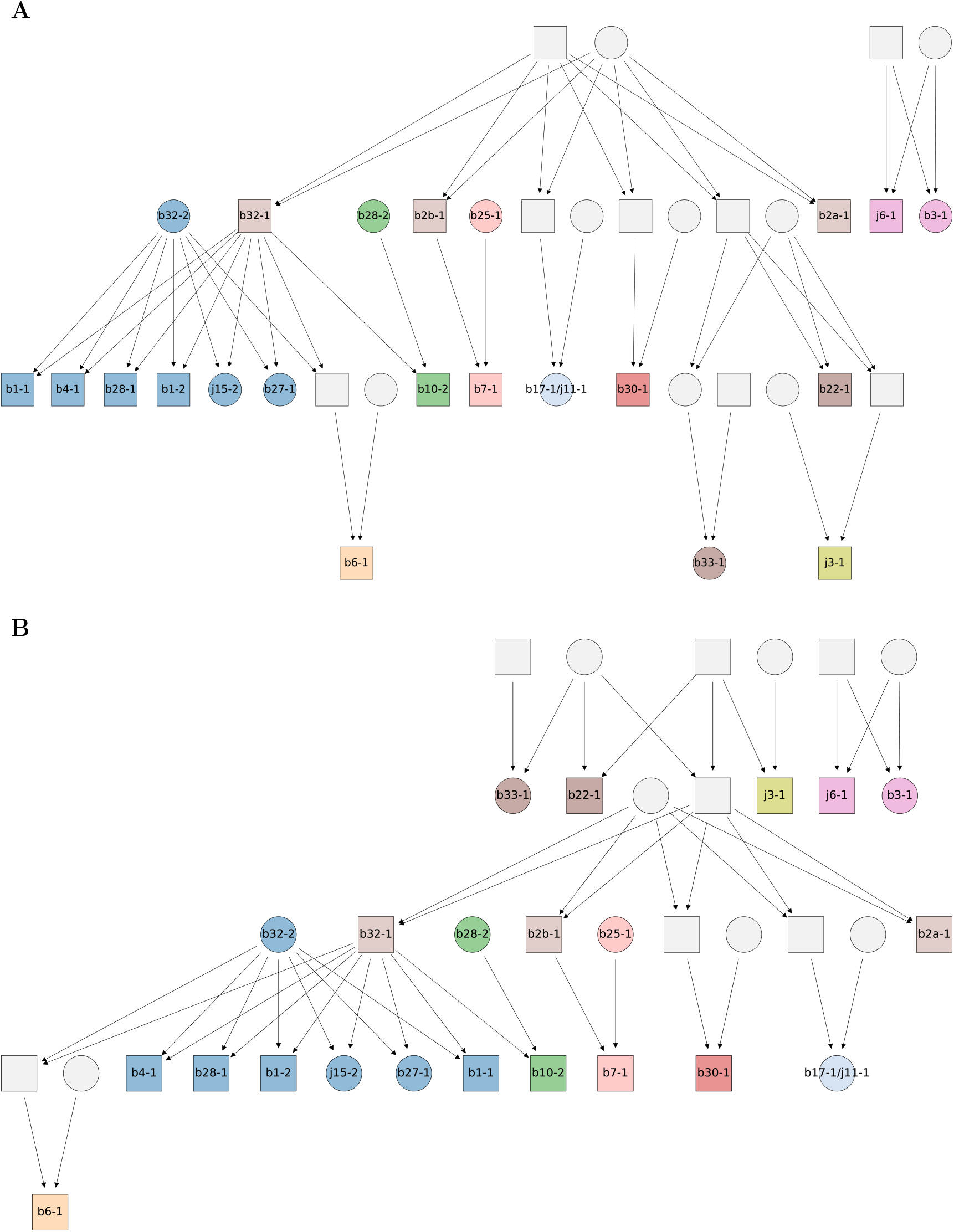
**(A)** The published Nepluyevsky pedigree [15] and **(B)** the original `repare`-reconstructed Nepluyevsky pedigree. Gray nodes without an ID label correspond to placeholder individuals. Nodes corresponding to non-placeholder individuals are colored by the individuals’ mitochondrial haplogroups. Unrelated individuals are not included in these plots.

**Figure 5.**
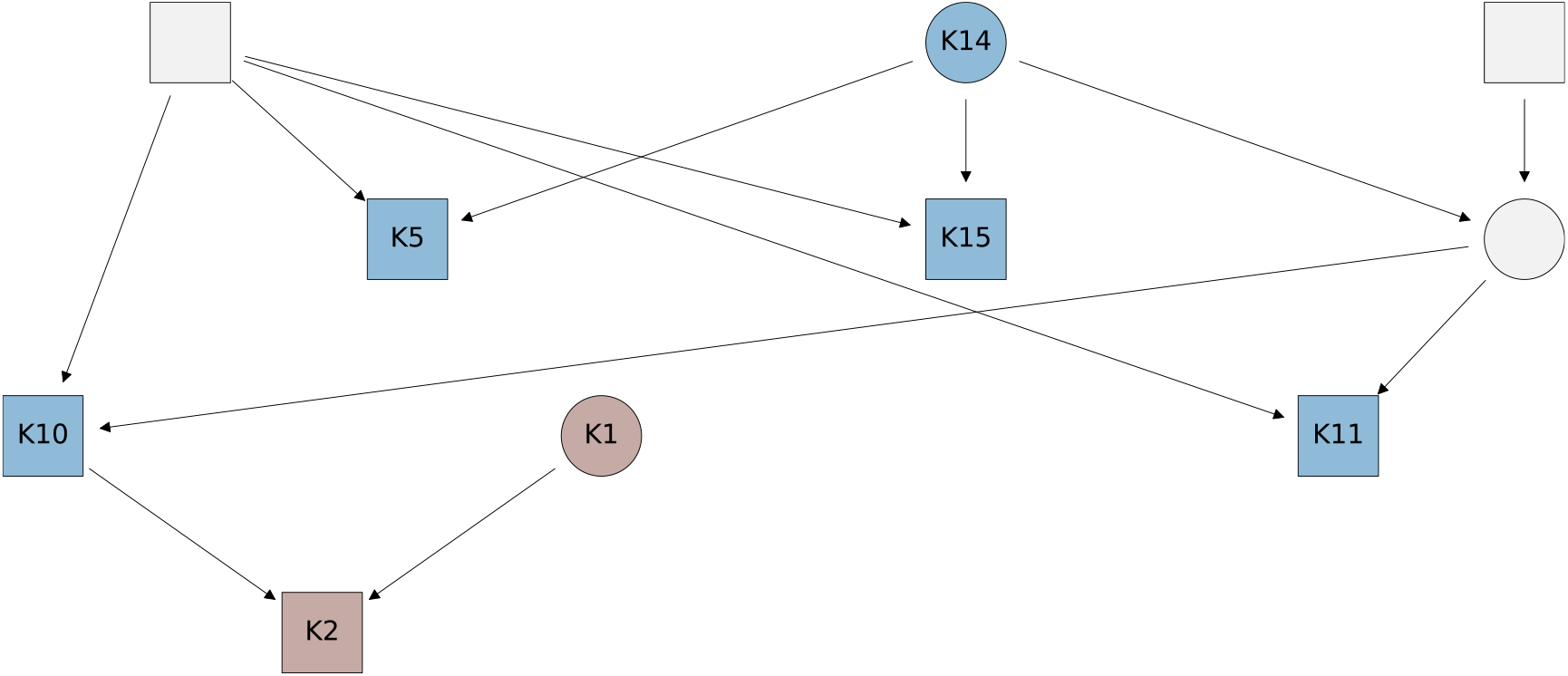
The `repare`-reconstructed Koszyce pedigree. Gray nodes without an ID label correspond to placeholder individuals. Nodes corresponding to non-placeholder individuals are colored by the individuals’ mitochondrial haplogroups.

#### 3.2.3 Koszyce site

To evaluate performance on a dataset including complex kinship relations, we use `repare` to reconstruct a Late Neolithic (sub)pedigree from the Koszyce site [21]. This site includes a pedigree inferred to contain complex kinship relations, including two pairs of individuals related at both the second and third degrees. This seven-individual pedigree, involving a subset of the Koszyce site’s related individuals, was reconstructed in a later analysis of the individuals’ kinship relations [31]. As such, we primarily use the kinship relations inferred by GRUPS-rs in this later analysis [31]. Since GRUPS-rs does not infer first-degree relation types, we use inferred first-degree relation types from the original article describing the Koszyce site [21]. We also obtain individual-level metadata, including haplogroups and skeletal age-at-death estimates, from the original article. Given this input information, `repare` successfully reconstructs the seven-individual pedigree. Notably, GRUPS-rs infers that K14 shares a first-degree relation with the mother of K10 and K11; it does not specify the exact relation type. `repare` successfully recovers this first-degree relation.

#### 3.2.4 Gurgy ‘les Noisats’ site

Finally, we assess `repare`’s reconstruction of a Neolithic pedigree from the Gurgy ‘les Noisats’ site [14]. Similar to the previous pedigrees, the published Gurgy pedigree was manually reconstructed using pairwise kinship relations and supporting information including haplogroups and skeletal age-at-death estimates. For the data used to reconstruct the published pedigree, degree-level kinship relations were inferred using `READ` [30], and first-degree relation types were inferred using `lcMLkin` [32]. In addition, uncertain relations were investigated using `BREADR` [35]. These data sources had many mutual conflicts, which were manually resolved by the pedigree authors. Therefore, instead of attempting to unify the multiple kinship data sources for input to `repare`, we evaluate reconstruction of the Gurgy pedigree using later-reported kinship data from `READv2` [27], a kinship inference method which infers both degree-level relations and first-degree relation types. With this input data, `repare` achieves a relation F1 score of 0.86 and a degree F1 score of 0.95 against the published pedigree.

Compared to the published Gurgy pedigree, the `repare`-reconstructed Gurgy pedigree contains the same number of first- and second-degree inconsistencies and fewer third-degree inconsistencies against the `READv2`-inferred relations. However, the `repare`-reconstructed Gurgy pedigree does not appear to be more plausible than the published Gurgy pedigree. The published Gurgy pedigree was validated using inferred IBD data, which offers a relatively independent measurement of a pedigree’s plausibility. We plot these published IBD data [14], inferred using `ancIBD` [12], for both the published pedigree and the `repare`-reconstructed pedigree Figure 6; the published pedigree is noticeably more concordant with the IBD data. In other words, although `repare` successfully reconstructs *most* of the Gurgy pedigree, its specific modifications to the input kinship relations appear to produce a pedigree that is overall less plausible than the published pedigree. As such, we believe this pedigree highlights a limitation of `repare`’s discrete inconsistency metric. `repare` treats all first- and second-degree inconsistencies as equivalent and all third-degree inconsistencies as equivalent. However, it is hypothetically possible to determine relative likelihoods of inconsistencies and assign more fine-grained scores to each inconsistency, for example using IBD data. It may also be possible to jointly infer likelihoods of entire pedigrees, bypassing the need to score individual inconsistencies. Ultimately, we believe the development of a continuous inconsistency scoring system, potentially leveraging additional sources of relatedness information such as IBD data, presents an opportunity for improvement in future versions of `repare`.

**Figure 6.**
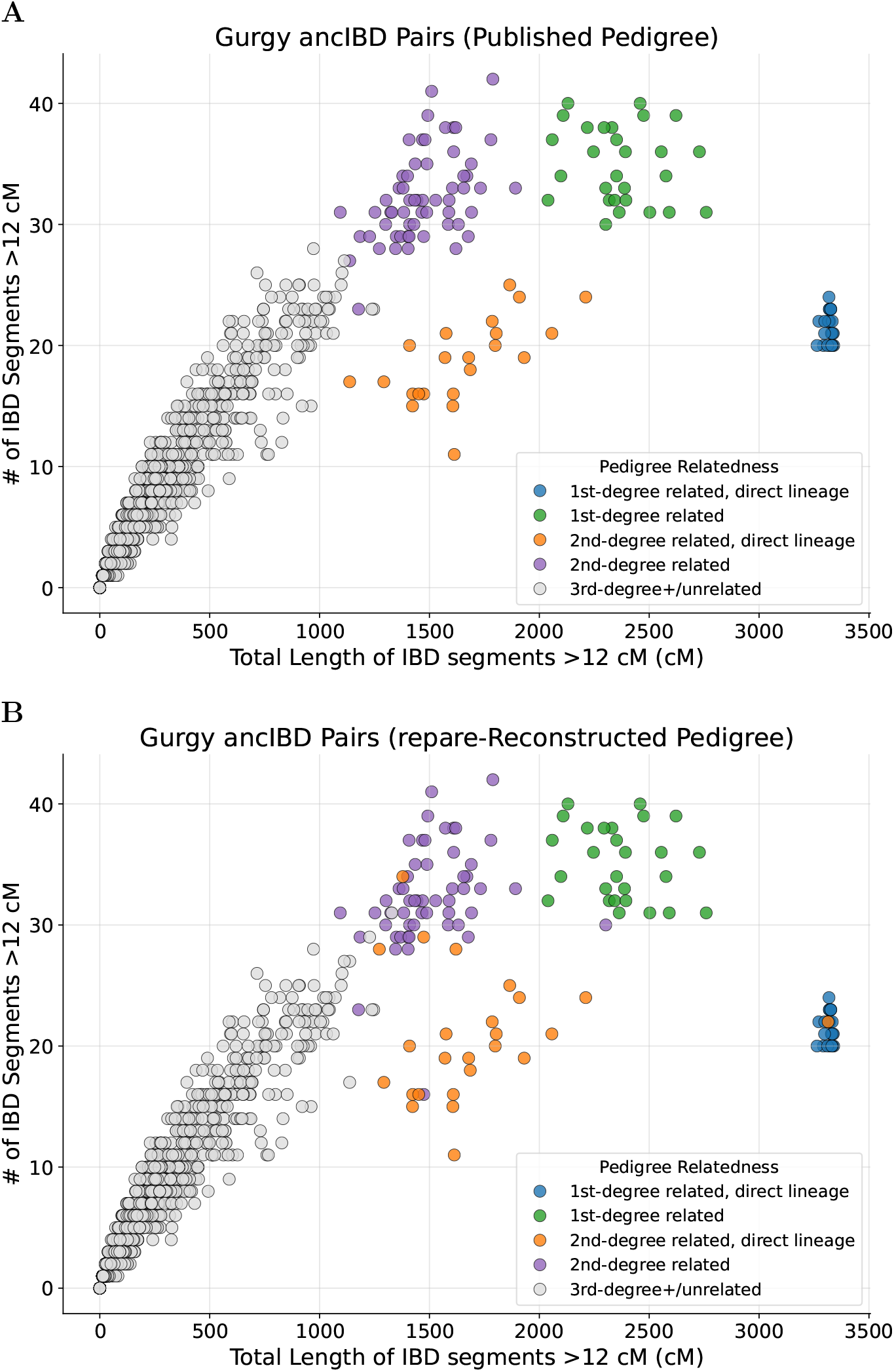
Inferred Gurgy IBD data for pairs of individuals, colored by kinship relations in **(A)** the published Gurgy pedigree [14] and **(B)** the `repare`-reconstructed Gurgy pedigree.

## 4 Discussion

We evaluate `repare` on a variety of simulated and published ancient pedigrees and find that it achieves strong reconstruction results under data quality conditions typical of aDNA. In addition, through reconstruction of published, manually reconstructed pedigrees, we highlight `repare`’s ability to flexibly integrate genetic and archaeological constraints alongside pairwise kinship relations (Results: Published pedigree reconstruction). In `repare`’s reconstructions of the Hazleton North and Nepluyevsky pedigrees, explicit incorporation of site-specific archaeological and genetic context, such as haplogroup rarities and recombination breakpoints, improved reconstruction to the point of successfully recovering the published pedigrees up to their first- and second-degree relations. `repare` is also able to reconstruct pedigrees involving complex relations, as we observe for the Koszyce seven-individual pedigree. We find that `repare` was unable to exactly recover the published Gurgy ‘les Noisats’ pedigree, possibly due to limitations in `repare`’s discrete inconsistency metric (Methods: Robustness to kinship inference errors). Improving the scoring of candidate pedigrees is therefore a promising avenue of future work.

As seen in `repare`’s performance on simulated pedigrees (Results: Simulated pedigree reconstruction), pedigree reconstruction is a process sensitive to input data quality. The exponential nature of iterative pedigree reconstruction makes it difficult to correct an excessive number of kinship inference errors. However, in sufficiently dense pedigrees, information from multiple adjacent kinship relations can help resolve pedigree topologies. In addition, it is still possible to make inferences about social dynamics and other first-order questions from pedigrees with some error. In this sense, reconstructing degree-level relations for an approximate pedigree topology can be nearly as productive as reconstructing exact relation types, which is a more difficult task.

To our knowledge, `repare` is the first pedigree reconstruction method to primarily utilize pairwise kinship relations. This makes `repare` particularly suitable for the reconstruction of ancient pedigrees. Existing ancient pedigrees reported in the literature are largely manually reconstructed, highlighting a need for automatic reconstruction methods that tolerate the relatively poor data quality of aDNA. We show that `repare` represents a powerful method for the automatic reconstruction of ancient pedigrees using only data commonly collected in aDNA analyses. In addition, `repare` can flexibly integrate user-inferred pedigree constraints, allowing users to guide reconstruction analyses through an iterative “human-in-the-loop” process.

## Acknowledgements

We thank Inñigo Olalde and Harald Ringbauer for insightful discussions on ancient pedigree reconstruction, and Egor Lappo for helpful advice on software reproducibility. This work was supported by a Director Discretionary Allocation from the Texas Advanced Computing Center (TACC) at The University of Texas at Austin and by the Allen Discovery Center program, a Paul G. Allen Frontiers Group advised program of the Paul G. Allen Family Foundation.

## Code availability

Code and documentation for `repare` are available at https://github.com/ehuangc/repare. The version of `repare` and associated data used in this work are archived on Zenodo [39].

## Data availability

No new DNA data were generated in this work. Data used to reconstruct ancient pedigrees are available from the supplementary materials of [13–15, 21].

## Contributions

E.C.H. developed the software, ran the experiments, and wrote the paper with input from all authors.

K.A.L. developed an initial version of the software. V.M.N. conceived and supervised the work.

## Ethics declarations

The authors report no conflicts of interest.

## References

[1] James Cussens et al. “Maximum Likelihood Pedigree Reconstruction Using Integer Linear Programming”. Genetic Epidemiology 37 (2013), pp. 69–83. doi: 10.1002/gepi.21686.

[2] Dan He et al. “IPED2: Inheritance Path Based Pedigree Reconstruction Algorithm for Complicated Pedigrees”. IEEE/ACM Transactions on Computational Biology and Bioinformatics 14 (2017), pp. 1094–1103. doi: 10.1109/TCBB.2017.2688439.

[3] Jisca Huisman. “Pedigree reconstruction from SNP data: parentage assignment, sibship clustering and beyond”. Molecular Ecology Resources 17 (2017), pp. 1009–1024. doi: 10.1111/1755-0998.12665.

[4] Ethan M. Jewett et al. “Bonsai: An efficient method for inferring large human pedigrees from genotype data”. The American Journal of Human Genetics 108 (2021), pp. 2052–2070. doi: 10.1016/j.ajhg.2021.09.013.

[5] Amy Ko and Rasmus Nielsen. “Composite likelihood method for inferring local pedigrees”. PLOS Genetics 13 (2017), e1006963. doi: 10.1371/journal.pgen.1006963.

[6] Markus Riester, Peter F. Stadler, and Konstantin Klemm. “FRANz: reconstruction of wild multi-generation pedigrees”. Bioinformatics 25 (2009), pp. 2134–2139. doi: 10.1093/bioinformatics/btp064.

[7] Doron Shem-Tov and Eran Halperin. “Historical Pedigree Reconstruction from Extant Populations Using PArtitioning of RElatives (PREPARE)”. PLOS Computational Biology 10 (2014), e1003610. doi: 10.1371/journal.pcbi.1003610.

[8] Jeffrey Staples et al. “PRIMUS: Rapid Reconstruction of Pedigrees from Genome-wide Estimates of Identity by Descent”. The American Journal of Human Genetics 95 (2014), pp. 553–564. doi: 10.1016/j.ajhg.2014.10.005.

[9] Swapan Mallick et al. “The Allen Ancient DNA Resource (AADR) a curated compendium of ancient human genomes”. Scientific Data 11 (2024), p. 182. doi: 10.1038/s41597-024-03031-7.

[10] Aurelien Ginolhac et al. “mapDamage: testing for damage patterns in ancient DNA sequences”. Bioinformatics 27 (2011), pp. 2153–2155. doi: 10.1093/bioinformatics/btr347.

[11] Jakob W. Sedig et al. “Combining ancient DNA and radiocarbon dating data to increase chronological accuracy”. Journal of Archaeological Science 133 (2021), p. 105452. doi: 10.1016/j.jas.2021.105452.

[12] Harald Ringbauer et al. “Accurate detection of identity-by-descent segments in human ancient DNA”. Nature Genetics 56 (2024), pp. 143–151. doi: 10.1038/s41588-023-01582-w.

[13] Chris Fowler et al. “A high-resolution picture of kinship practices in an Early Neolithic tomb”. Nature 601 (2022), pp. 584–587. doi: 10.1038/s41586-021-04241-4.

[14] Maïtë Rivollat et al. “Extensive pedigrees reveal the social organization of a Neolithic community”. Nature 620 (2023), pp. 600–606. doi: 10.1038/s41586-023-06350-8.

[15] Jens Blöcher et al. “Descent, marriage, and residence practices of a 3,800-year-old pastoral community in Central Eurasia”. Proceedings of the National Academy of Sciences 120 (2023), e2303574120. doi: 10.1073/pnas.2303574120.

[16] Maciej Chyleński et al. “Patrilocality and hunter-gatherer-related ancestry of populations in East-Central Europe during the Middle Bronze Age”. Nature Communications 14 (2023), p. 4395. doi: 10.1038/s41467-023-40072-9.

[17] Guido Alberto Gnecchi-Ruscone et al. “Network of large pedigrees reveals social practices of Avar communities”. Nature 629 (2024), pp. 376–383. doi: 10.1038/s41586-024-07312-4.

[18] Alissa Mittnik et al. “Kinship-based social inequality in Bronze Age Europe”. Science 366 (2019), pp. 731–734. doi: 10.1126/science.aax6219.

[19] Sandra Penske et al. “Kinship practices at the early bronze age site of Leubingen in Central Germany”. Scientific Reports 14 (2024), p. 3871. doi: 10.1038/s41598-024-54462-6.

[20] Ricardo Rodríguez-Varela et al. “Five centuries of consanguinity, isolation, health, and conflict in Las Gobas: A Northern Medieval Iberian necropolis”. Science Advances 10 (2024), eadp8625. doi: 10.1126/sciadv.adp8625.

[21] Hannes Schroeder et al. “Unraveling ancestry, kinship, and violence in a Late Neolithic mass grave”. Proceedings of the National Academy of Sciences 116 (2019), pp. 10705–10710. doi: 10.1073/pnas.1820210116.

[22] Frederik Valeur Seersholm et al. “Repeated plague infections across six generations of Neolithic Farmers”. Nature 632 (2024), pp. 114–121. doi: 10.1038/s41586-024-07651-2.

[23] Anna Szëcsënyi-Nagy et al. “Ancient DNA reveals diverse community organizations in the 5th millennium BCE Carpathian Basin”. Nature Communications 16 (2025), p. 5318. doi: 10.1038/s41467-025-60368-2.

[24] Jincheng Wang et al. “Genomic profiling of a six-generation patrilineal family of the Ming-Qing dynasties in China”. iScience 28 (2025), p. 112968. doi: 10.1016/j.isci.2025.112968.

[25] Ke Wang et al. “Ancient DNA reveals reproductive barrier despite shared Avar-period culture”. Nature 638 (2025), pp. 1007–1014. doi: 10.1038/s41586-024-08418-5.

[26] Eren Yüncü et al. “Female lineages and changing kinship patterns in Neolithic Catalhöyük”. Science 388 (2025), eadr2915. doi: 10.1126/science.adr2915.

[27] Erkin Alacamli et al. “READv2: advanced and user-friendly detection of biological relatedness in archaeogenomics”. Genome Biology 25 (2024), p. 216. doi: 10.1186/s13059-024-03350-3.

[28] Daniel M. Fernandes et al. “TKGWV2: an ancient DNA relatedness pipeline for ultra-low coverage whole genome shotgun data”. Scientific Reports 11 (2021), p. 21262. doi: 10.1038/s41598-021-00581-3.

[29] Kristian Hanghøj et al. “Fast and accurate relatedness estimation from high-throughput sequencing data in the presence of inbreeding”. GigaScience 8 (2019), giz034. doi: 10.1093/gigascience/giz034.

[30] Jose Manuel Monroy Kuhn, Mattias Jakobsson, and Torsten Günther. “Estimating genetic kin relationships in prehistoric populations”. PLOS ONE 13 (2018), e0195491. doi: 10.1371/journal.pone.0195491.

[31] Maël Lefeuvre et al. “GRUPS-rs, a high-performance ancient DNA genetic relatedness estimation software relying on pedigree simulations”. Human Population Genetics and Genomics 4 (2024). doi: 10.47248/hpgg2404010001.

[32] Mikhail Lipatov et al. Maximum Likelihood Estimation of Biological Relatedness from Low Coverage Sequencing Data. bioRxiv. 2015. doi: 10.1101/023374.

[33] Emil Nyerki et al. “correctKin: an optimized method to infer relatedness up to the 4th degree from low-coverage ancient human genomes”. Genome Biology 24 (2023), p. 38. doi: 10.1186/s13059-023-02882-4.

[34] Divyaratan Popli, Stëphane Peyrëgne, and Benjamin M. Peter. “KIN: a method to infer relatedness from low-coverage ancient DNA”. Genome Biology 24 (2023), p. 10. doi: 10.1186/s13059-023-02847-7.

[35] Adam B. Rohrlach et al. “BREADR: An R Package for the Bayesian Estimation of Genetic Relatedness from Low-coverage Genotype Data”. Journal of Open Source Software 10 (2025), p. 7916. doi: 10.21105/joss.07916.

[36] Ryan K. Waples, Anders Albrechtsen, and Ida Moltke. “Allele frequency-free inference of close familial relationships from genotypes or low-depth sequencing data”. Molecular Ecology 28 (2019), pp. 35–48. doi: 10.1111/mec.14954.

[37] Richard S. Sutton and Andrew Barto. Reinforcement Learning: An Introduction. Second edition. Adaptive computation and machine learning. Cambridge, Massachusetts London, England: The MIT Press, 2020.

[38] Maël Lefeuvre, Marie-Claude Marsolier, and Cëline Bon. BADGER: evaluating the performance of ancient DNA genetic relatedness estimation methods using high-fidelity pedigree simulations. Research Square. 2025. doi: 10.21203/rs.3.rs-7045281/v1.

[39] Edward C. Huang, Kevin A. Li, and Vagheesh M. Narasimhan. repare. Version v0.1.4. 2026. doi: 10.5281/ZENODO.18364224.

