## Supplementary Information for "Fault-tolerant pedigree reconstruction from pairwise kinship relations"

### Contents

Supplementary Note 1: Ancient pedigree simulation pg. S1

Supplementary Note 2: Analysis of simulated pedigree reconstruction pg. S2

Supplementary Note 3: Algorithm runtime and memory usage pg. S6

### Supplementary Note 1: Ancient pedigree simulation

To simulate ancient pedigrees, our custom simulator first generates four-generation pedigrees under simple demographic rules. The simulator starts with a small founder cohort of three individuals; at least two of these individuals reproduce with each other. Then, the simulator iterates through each individual, generation by generation, assigning mates and generating offspring. Each new individual is assigned a genetic sex and relevant haplogroup(s), as well as a binary ground-truth "can_have_children" status. All individuals with a true ground-truth "can_have_children" status will produce offspring. Offspring are assigned the appropriate haplogroups of their parents, and non-fixed haplogroups are drawn from a small discrete pool. If an individual can have children, the simulator assigns a mate for this target individual by either generating a new individual or randomly sampling an existing individual who can have children and is within one generation of the target individual. We select mates from existing individuals with some probability in order to promote consanguineous mating. The values of the pedigree simulator's demographic parameters were determined *a posteriori* based on the general characteristics of the resulting pedigrees.

After generating a pedigree, the simulator masks and corrupts its data to approximate real-world inputs to `repare`. First, since pedigrees are virtually never completely sampled, individuals in the pedigree are randomly removed from `repare`'s input data. For kept individuals, the simulator consistently preserves genetic sex and haplogroup information from the simulated pedigree but only preserves some individuals whose ground-truth "can_have_children" status is false, since skeletal age-at-death is only one reason why individuals might not have reproduced. Then, to simulate errors in kinship relation inference, ground-truth kinship relations and first-degree relation types are corrupted in `repare`'s input data. We do not simulate any relation type inferences for relations beyond the first degree or for non-first-degree relations that have been corrupted to the first degree. In addition, when a pair of individuals shares multiple kinship relations, for example due to inbreeding, we only use the closest kinship relation (before relation corruption) as input for pedigree reconstruction; this is to further approximate the data generated by kinship inference software. We ensure that at least two individuals and one inferred first- or second-degree kinship relation are present in the resulting kinship relation dataset.

We derive our simulator's kinship relation corruption rates from `KIN`'s results in the article describing `BADGER`, a benchmarking pipeline for ancient kinship inference [1]. To approximate kinship inference performance at varying input data qualities, we use `KIN`'s relation classification results at simulated sequence coverages of 0.1x, 0.2x, 0.5x, and 1.0x. For each coverage level, we assume a 0.01 error rate in first-degree relation type inference, which is conservatively estimated from the results of the article describing `KIN` [2].

### Supplementary Note 2: Analysis of simulated pedigree reconstruction

We report summary statistics of the 100 simulated ground-truth pedigrees used in our experiments (Results: Simulated pedigree reconstruction) (Figure S1). Since `repare` considers kinship relations up to the third degree, we define inbred individuals as those with parents related at the third degree or closer. Note that relation and degree F1 scores are computed over first- and second-degree relations (Methods: Pedigree reconstruction).

To investigate the effect of pedigree size and inbreeding on `repare`'s performance, we also report pedigree-level reconstruction results from two simulated pedigree experiments (Methods: Simulated pedigree reconstruction) (Figure S2). When $p(\text{mask node}) = 0.4$ and simulated sequence coverage is 0.5x, we find that `repare`'s reconstruction performance appears robust to changes in ground-truth pedigree size. We also observe a potential slightly negative relationship between ground-truth inbreeding proportion and reconstruction performance. In summary, at a simulated sequence coverage of 0.5x, we find that `repare` remains robust to increases in pedigree size and, potentially to a lesser degree, increases in inbreeding proportion. Holding $p(\text{mask node})$ at 0.4, we also report pedigree-level reconstruction results at a simulated sequence coverage of 0.1x. At this lower simulated sequence coverage, we note that `repare`'s reconstruction performance appears less robust to increases in pedigree size. We also report the distribution of pedigree-level results for these two simulated pedigree experiments (Figure S3).

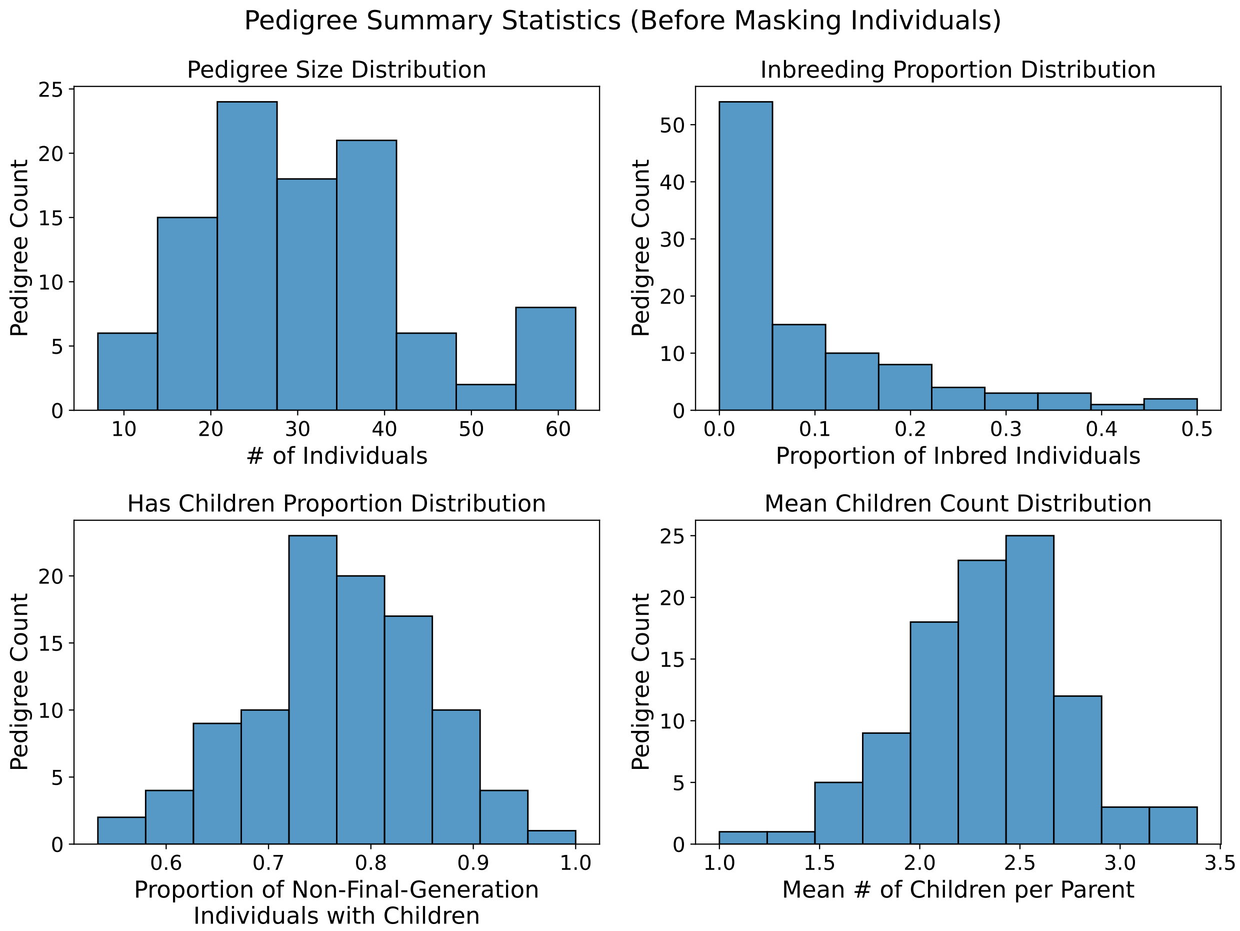


Figure S1: Summary statistics of the 100 simulated pedigrees used to test `repare`'s reconstruction performance.

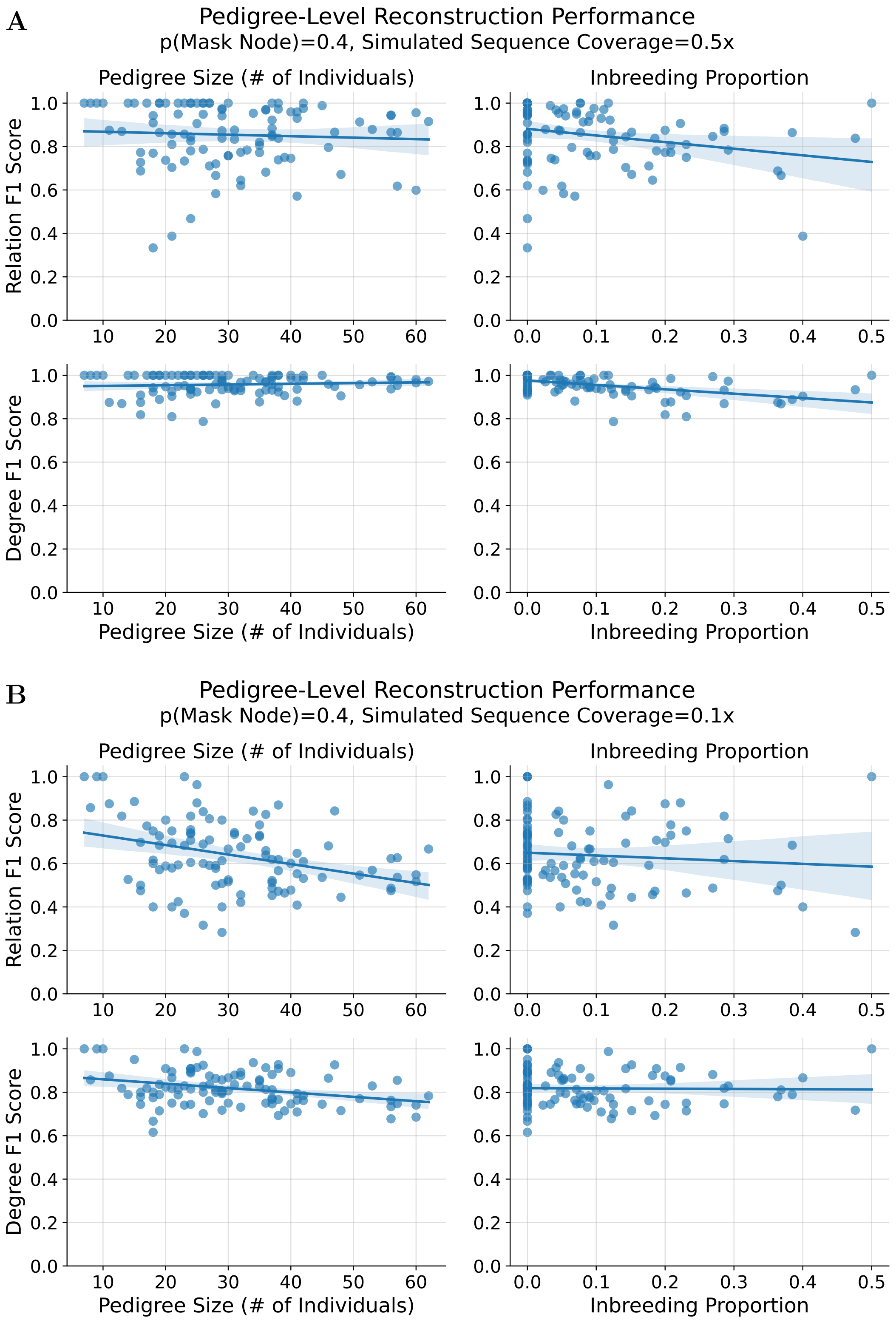


Figure S2: Pedigree-level reconstruction results for simulated pedigrees with **(A)** $p(\text{mask node}) = 0.4$ and simulated sequence coverage of 0.5x and **(B)** $p(\text{mask node}) = 0.4$ and simulated sequence coverage of 0.1x. Each point corresponds to one simulated pedigree and its reconstruction result. Pedigree sizes and inbreeding proportions are computed using unmasked, ground-truth simulated pedigrees. Regression lines are computed with ordinary least-squares and plotted with 95% bootstrap confidence intervals.

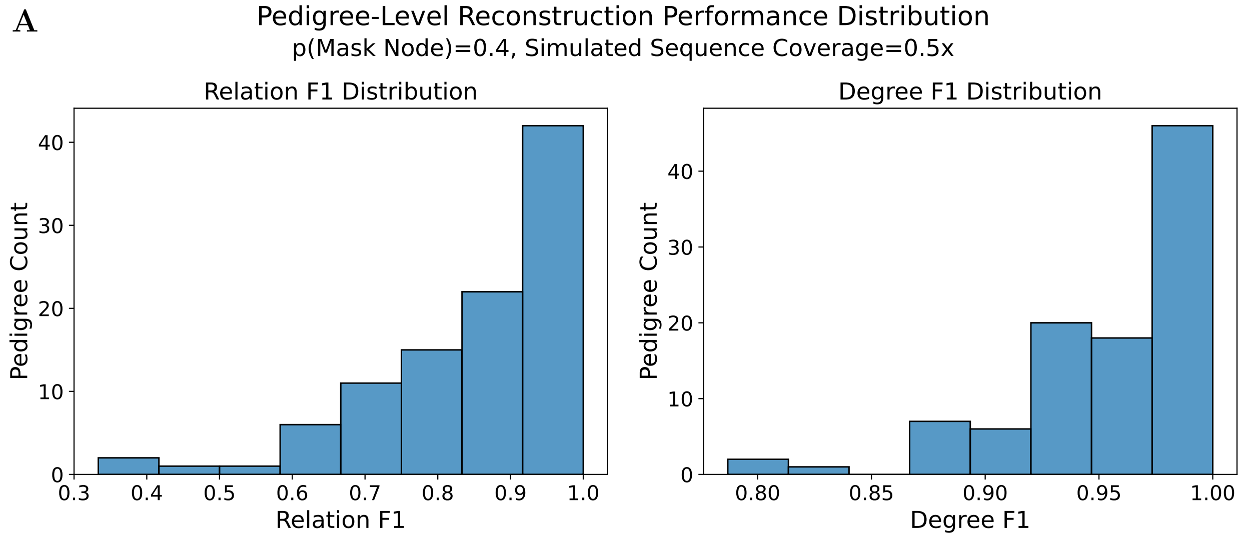


Figure S3: Distributions of pedigree-level reconstruction results for simulated pedigrees with **(A)** $p(\text{mask node}) = 0.4$ and simulated sequence coverage of 0.5x and **(B)** $p(\text{mask node}) = 0.4$ and simulated sequence coverage of 0.1x.

### Supplementary Note 3: Algorithm runtime and memory usage

There are six second-degree relation types: aunt/uncle-nephew/niece, nephew/niece-aunt/uncle, grandparent-grandchild, grandchild-grandparent, half-siblings, and double first cousins. After considering the genetic sex of the intermediary individual (e.g., maternal vs. paternal grandparent) and the two possible configurations of double first cousins (same-genetic-sex siblings vs. cross-genetic-sex siblings), this yields 12 possible second-degree relation types. Additionally, `repare` also considers inferred second-degree relations as potential first-degree relations (parent-child, child-parent, or siblings) and potential false positives (unrelated); this means there are 16 possible results of incorporating a second-degree relation. Therefore, since `repare` explicitly incorporates up to second-degree relations, the worst-case number of potential pedigrees corresponding to a set of degree-level kinship relations is $O(16^n)$, where $n$ is the number of first- and second-degree input kinship relations.

Exponential growth of the candidate pedigree set in the number of input relations is extremely undesirable, especially since the number of input relations itself can increase quadratically with the number of analyzed individuals. Downsampling the set of candidate pedigrees to a fixed size after each algorithm iteration allows us to constrain the number of pedigrees enumerated per iteration. After each iteration, we keep at most $k$ candidate pedigrees, where $k$ is a user-defined constant that defaults to 1000. During each iteration, since we incorporate one input kinship relation, we generate at most $16 \cdot k$ candidate pedigrees before downsampling back to $k$ candidate pedigrees. Therefore, through this reconstruction process, the maximum number of pedigrees enumerated is $O(16kn) = O(n)$, where $k$ is a constant that defaults to 1000 and $n$ is the number of input kinship relations.

Although the maximum number of enumerated pedigrees is linear in the number of input relations, this does not imply that algorithm *runtime* is linear in the number of input relations. For example, in each algorithm iteration, `repare` must validate each generated candidate pedigree and count inconsistencies, including for third-degree relations, for each retained candidate pedigree; these actions themselves are dependent in various ways on the number of input relations and candidate pedigree size (Methods: Pedigree pruning) (Methods: Robustness to kinship inference errors). As such, an exact runtime bound is impractical to determine, and we instead report an empirical benchmark of `repare`'s runtime and memory usage (Figure S4). To generate inputs to `repare`, we randomly sample sets of individuals from the Gurgy dataset [3], subsetting individual-level and relation-level input data to include only these individuals. Then, we run `repare` three times for each version of the input data, recording the runtime and peak resident set size (RSS) of each run. Peak RSS measures the largest amount of non-swapped physical memory used by the process at any point during execution. We observe that both runtime and peak RSS increase reasonably with the number of input first- and second-degree relations. This benchmark was run on a Frontera Cascade Lake compute node (dual-socket Intel Xeon Platinum 8280) [4].

**A**

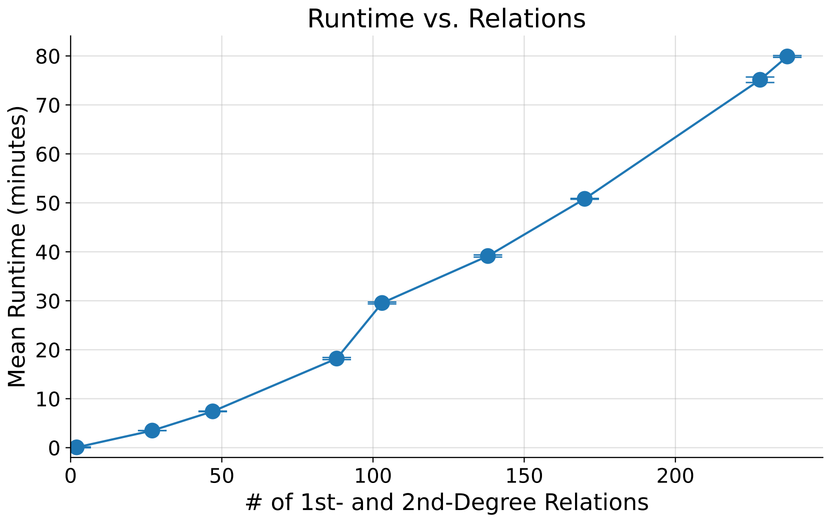


**B**

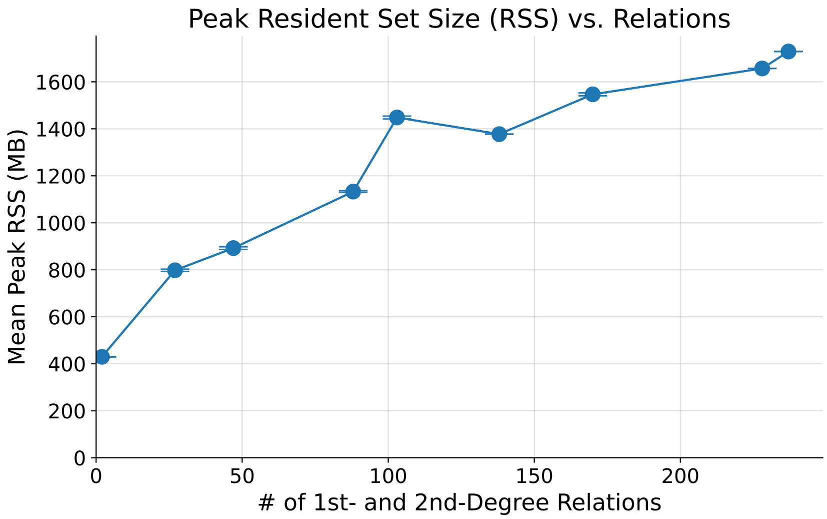


Figure S4: **(A)** Runtime and **(B)** peak resident set size (RSS) of `repare`, plotted against the number of input first- and second-degree relations. Each point represents the mean result of three replicate runs, and error bars represent standard deviations.

### References

[1] Maël Lefeuvre, Marie-Claude Marsolier, and Céline Bon. *BADGER: evaluating the performance of ancient DNA genetic relatedness estimation methods using high-fidelity pedigree simulations*. Research Square. 2025. DOI: 10.21203/rs.3.rs-7045281/v1.

[2] Divyaratan Popli, Stéphane Peyrégne, and Benjamin M. Peter. "KIN: a method to infer relatedness from low-coverage ancient DNA". *Genome Biology* 24 (2023), p. 10. DOI: 10.1186/s13059-023-02847-7.

[3] Maïté Rivollat et al. "Extensive pedigrees reveal the social organization of a Neolithic community". *Nature* 620 (2023), pp. 600–606. DOI: 10.1038/s41586-023-06350-8.

[4] *Frontera - TACC HPC Documentation.* URL: https://docs.tacc.utexas.edu/hpc/frontera/ (visited on 01/19/2026).
